# Compromised actin dynamics underlie the orofacial cleft in Baraitser-Winter Cerebrofrontofacial Syndrome with a variant in *ACTB*

**DOI:** 10.1101/2024.04.04.587685

**Authors:** Takayuki Tsujimoto, Yushi Ou, Makoto Suzuki, Yuka Murata, Toshihiro Inubushi, Miho Nagata, Yasuki Ishihara, Ayumi Yonei, Yohei Miyashita, Yoshihiro Asano, Norio Sakai, Yasushi Sakata, Hajime Ogino, Takashi Yamashiro, Hiroshi Kurosaka

**Author notes:** Corresponding Author: Hiroshi Kurosaka, Department of Orthodontics and Dentofacial, Orthopedics, Osaka University Graduate School of Dentistry, 1-8 Yamadaoka, Suita, Osaka, 565-0871, Japan. **Conflict of interest statement** The authors have declared that no conflict of interest exists.

## Abstract

Craniofacial anomalies encompassing the orofacial cleft are associated with >30% of systemic congenital malformations. Baraitser-Winter Cerebrofrontofacial syndrome (BWCFF) is a rare genetic disorder attributed to variants in the actin beta (*ACTB*) or actin gamma genes that are correlated with a range of craniofacial abnormalities, including cleft lip and/or palate. The underlying pathological mechanism of BWCFF remains elusive, and it is necessary to investigate the etiology of orofacial clefts in BWCFF.

In this study, we identified a missense variant (c.1043C>T: p.S348L) in the *ACTB* gene from a patient with BWCFF and concomitant cleft lip and palate. Furthermore, we performed functional assessments of this variant using various disease models such as the MDCK cell line and *Xenopus laevis*. These models revealed a compromised capacity of mutated ACTB to localize at the epithelial junction, consequently affecting the behavior of epithelial cells. Additionally, we made the noteworthy discovery that mutated ACTB exhibited an impaired ability to bind PROFILIN1, a critical factor in actin polymerization. This defective ability may contribute to the molecular etiology of aberrant epithelial cell adhesion and migration, resulting in orofacial cleft formation in BWCFF.

**Author Summary:** Advances in clinical sequencing have rapidly elucidated the genetic basis of rare diseases affecting multiple organs. In this study, we investigate the role of a variant in the *ACTB* gene, which is fundamental to cell structure, in causing orofacial clefts in the Baraitser-Winter Cerebrofrontofacial Syndrome (BWCFF), known for its distinctive facial anomalies. Using epithelial cell lines and frog embryo models, we investigated the effects of the *ACTB* gene alteration on cellular functions. Our results indicate that this genetic change disrupts the normal adhesion and movement of epithelial cells, processes that are critical for facial development. This research underscores the importance of the *ACTB* gene in shaping the face and suggests new directions for future research, potentially informing the diagnosis and treatment of BWCFF and related conditions.

## Introduction

Baraitser-Winter Cerebrofrontofacial syndrome (BWCFF) is a congenital disorder characterized by diverse phenotypes, including short stature, ptosis, lissencephaly, and characteristic craniofacial features, such as hypertelorism and occasional orofacial cleft[1, 2]. Actin beta (ACTB), a known causative gene of BWCFF, encodes an actin protein that forms actin filaments that comprise the cytoskeleton[3]. Based on this background, the phenotypes in a variety of organs in BWCFF are thought to be caused by malfunctioning actin complexes.

Actin proteins have been reported to perform a diverse range of functions by binding to and forming complexes with several proteins[4]. Typical functions include cell migration and adhesion through homeostatic behaviors, including actin polymerization and depolymerization of actin-associated proteins[5, 6]. Cell migration commences with adhesion to the extracellular matrix at the tips of the extended filopodia and lamellipodia, which serve as fulcrums for the repeated traction and detachment of the cell body in the direction of movement[7]. Actin filaments also localize at intercellular junctions and play critical roles in cell adhesion by forming adherence junctions via different classes of Cadherin proteins[5].

A distinctive craniofacial phenotype of BWCFF is an orofacial cleft that occurs in approximately 20% of the cases[1]. Orofacial cleft is the most common congenital disorder in the oral and maxillofacial regions, and developmental defects at any step of palatogenesis play a major role in its pathogenesis. Secondary palate formation begins with the elongation of secondary palatal processes at approximately six weeks of gestation in humans. The right and left secondary palatine processes, which originate from the maxillary process, grow vertically on the lateral surface of the tongue, move upward with the extension of the processes, and eventually come into contact with each other at the midline to initiate fusion. At this stage, the midline edge epithelial seam (MES), an epithelial layer derived from the palatine processes, remains in the area of palatine process fusion[8]. One of the critical steps in the fusion of facial processes is MES removal to achieve mesenchymal continuity[8]. The mechanisms by which MES disappears have been extensively studied, including epithelial cell migration[9], apoptosis[10], epithelial-mesenchymal transformation (EMT)[11], and cell protrusion[12]. Interestingly, several studies have revealed the importance of actin polymerization (F-actin), with the manifestation of orofacial cleft phenotype when its activity is inhibited.

Given the genetic etiology and orofacial cleft phenotype in BWCFF, there is a compelling indication that aberrations in the actin complex during embryonic development result in orofacial cleft. In recent years, new variants have been reported in BWCFF; however, detailed functional analyses of each variant have not been performed. In particular, the mechanism by which *ACTB* variants cause orofacial clefts is largely unknown. In this study, we identified a pathological missense variant in *ACTB* (c.1043C > T: p.S348L) in a patient with BWCFF and cleft lip and palate. We further utilized multiple disease models to elucidate the molecular and cellular etiology of orofacial clefts in conjunction with malfunctioning actin complexes.

## Results

### Missense variant in actin beta *(ACTB)* (c.1043C > T:p.S348L) result in Baraitser-Winter Cerebrofrontofacial syndrome (BWCFF) with multiple craniofacial defects

A 9-year-old boy presented with systemic symptoms such as bilateral cleft lip (Figure1A) and complete cleft palate (Figure1B), thin calvarial bone (Figure1C), ptosis, hearing impairment, otitis media, hyperterolism, ventricular septal defect, atrial septal defect, small intestinal stenosis, cryptorchidism, and intellectual disability. He remained undiagnosed and underwent trio exome analysis along with his unaffected parents.

Subsequently, we identified a missense variant in the final exon of the *ACTB* gene (c.1043C>T: p.S348L), which led to a genetic diagnosis of BWCFF (Figure 1A–D). This variant was located in exon 5, and has been previously reported to cause BWCFF [13, 14] (Figure 1E).

**Figure 1.**
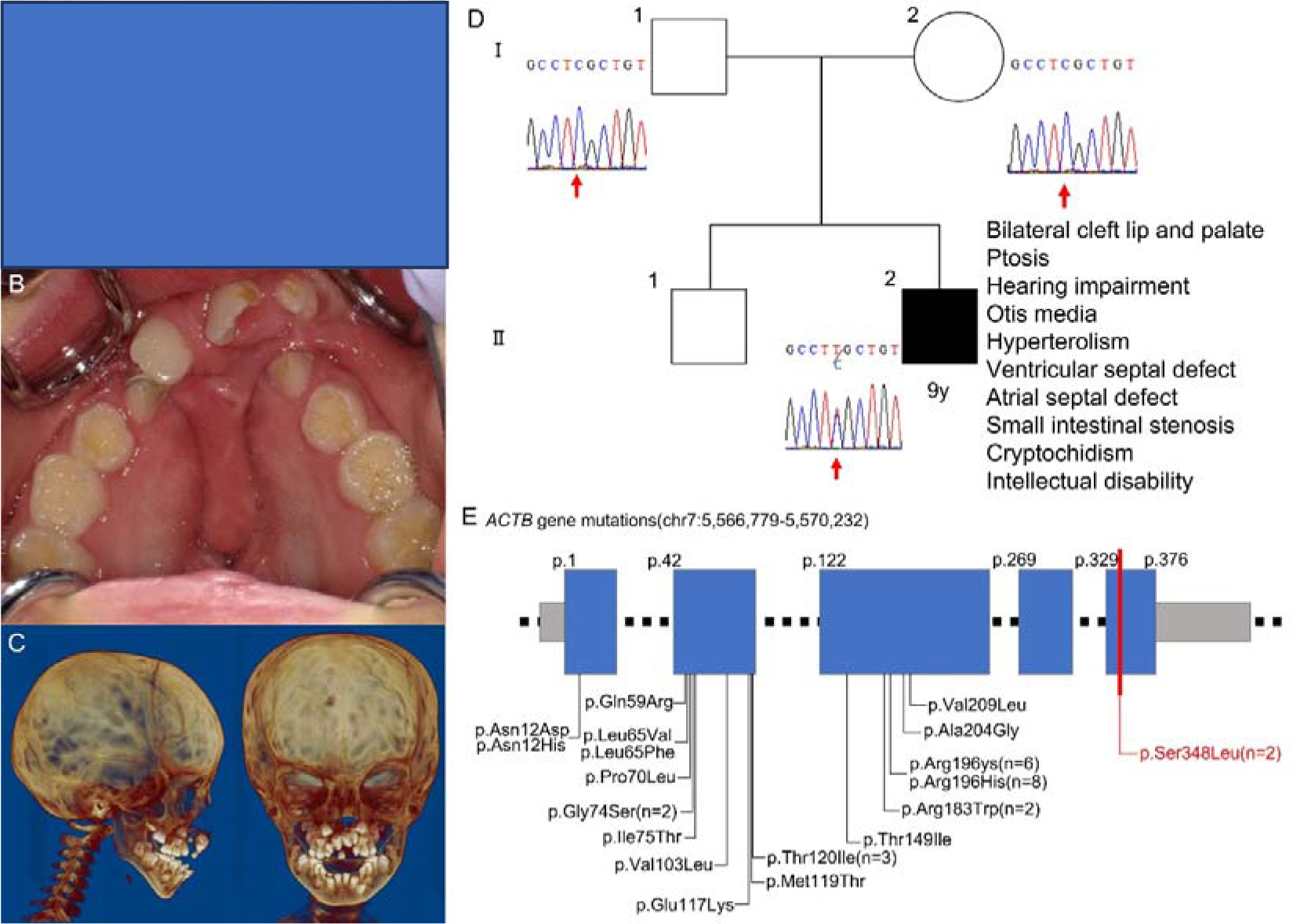
Craniofacial appearance and Identification of the missence variant in the actin beta (*ACTB)* (A) Facial, (B) intraoral, and (C) Computed Tomography imaged of the patient at the age of 6. (D) The pedigree of the present case with chromatogram and major phenotypes. (E) Summary of reported variants of *ACTB* in Baraitser-Winter Cerebrofrontofacial syndrome (BWCFF). A missense variant detected in present study is shown in red at the final exon of the *ACTB* gene (c.1043C > T: p.S348L).

### Actin beta *(ACTB)* expression in embryonic frontonasal process and fusing secondary palatal shelves

*In situ* hybridization of E9.5 mouse embryo demonstrated widespread expression of *ACTB* mRNA throughout the body, with certain tissues exhibiting higher expression, such as the frontonasal process and first branchial arch (Figure 2A). Notably, as embryonic development progressed, an increase in the contrast of expression was observed, with greater intensity in the frontonasal processes, branchial arches, and limb buds in the E10.5 embryo (Figure 2B). Notably, strong signal of *Actb* was detected in the epithelium of growing palatal shelves and midline edge epithelial seam (MES) at the fused secondary palatal shelves (Figure 2C and D). Furthermore, immunohistochemistry of ACTB together with the epithelial marker E-cadherin revealed a strong ACTB signal in the epithelial seam of the secondary palate (Figure 2E–H, Supplemental figure 1). Neither negative control of *in situ* hybridization and immunohistochemistry showed detectable signal (Supplemental figure 2). It is also important to note that negative control both *in situ* hybridization and immunohistochemistry did not show detectable signal of *Actb* (Supplemental figure 2). However, it is essential to note that the signal was ubiquitously detected throughout various tissues, including oral epithelial and mesenchymal cells. This is attributable to its function as a housekeeping gene across diverse tissues, with the level of expression subject to context-dependent alterations.

**Figure 2.**
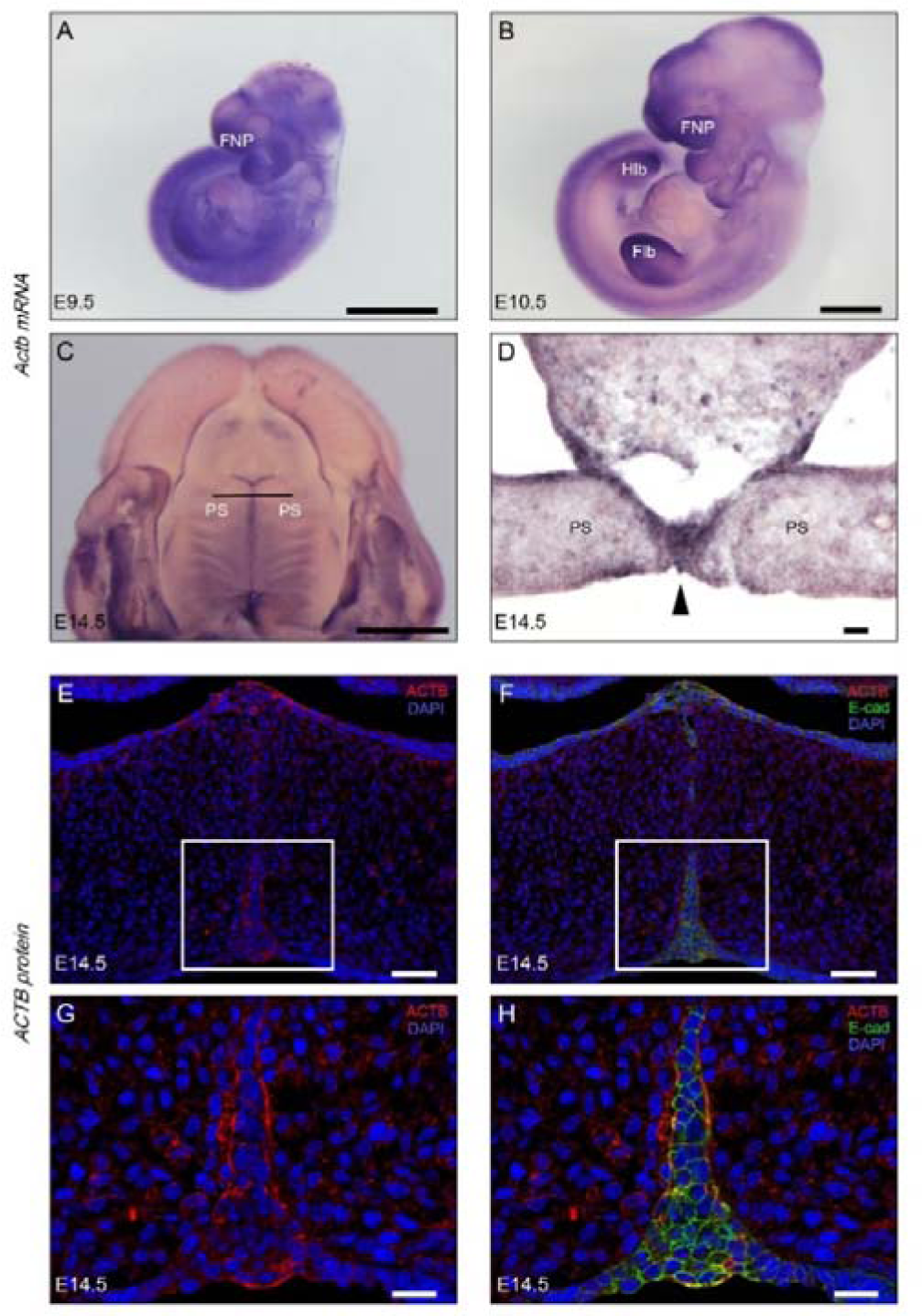
Expression pattern of actin beta (*Actb*) in embryonic craniofacial development. (A–D) *Actb* expression in different stage and tissue during mice embryonic development. Whole mount *in situ* hybridization of *Actb* using E9.5 (A) and E10.5 (B) mice embryo. (C) Ventral view of dissected maxilla at E14.5. Frontal section of E14.5 maxilla at the location of black line shown in C. Black arrowheads indicate midline edge epithelial seam (MES). (E-H) Immunohistochemistry of ACTB (red) and E-cadherin (green) of frontal section of embryonic head of E14.5. (G and H) show magnified images of the white squares of panels E and F, respectively. FNP, frontal nasal process. Flb, forelimb bud. HIb, hind limb bud. PS, palatal shelf. Scale bar, 1 mm (A–D), 50 μm (E, F), and 20 μm (G, H).

### Overexpression of mutant ACTB (p.S348L) in MDCK cells affects cell migration

One of the major mechanisms for preventing the secondary palate to fuse is retarded MES cell migration[9]. Therefore, we used MDCK cells overexpressing GFP-labeled ACTB (p. S348L) in the migration assay. We observed that the mutant ACTB (p.S348L)-overexpressing cell group displayed slower migration over time when compared with the wild-type ACTB overexpressing cell group (Figure 3A–D). Furthermore, the area of cell migration was measured over time by repeating the experiment thrice, revealing that the population of mutant ACTB (p.S348L)-overexpressing cells had predominantly smaller areas filled with cell migration 7 h after the beginning of cell culture which eventually close the wound after that (Supplemental figure 3A). These findings indicate that the overexpression of mutant ACTB (p.S348L) adversely affects epithelial cell migration.

**Figure 3.**
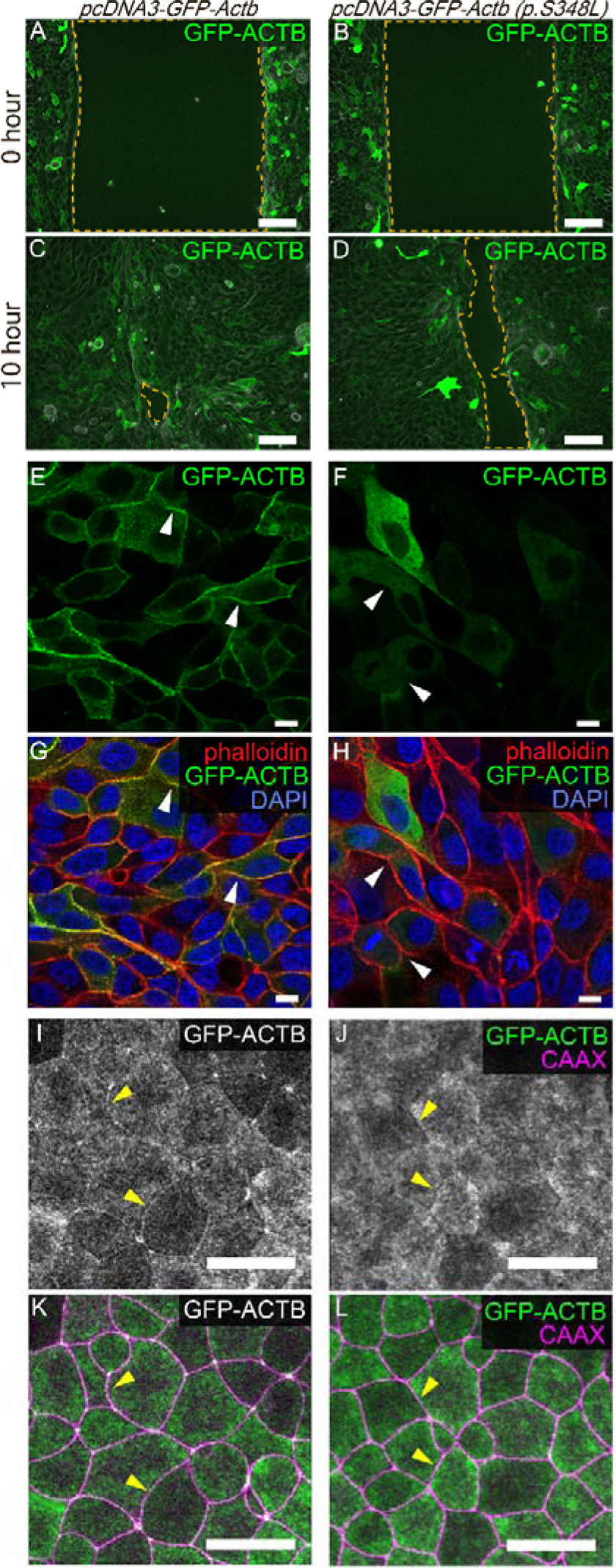
Functional assays of MDCK cells with overexpressed ACTB. (A–D) Time course of wound healing assay using MDCK cells with overexpressed GFP labeled ACTB. (A) Immediately after beginning the assay with wild type and (B) mutant ACTB (p.S348L). (C) 10 hours after beginning the assay with wild type and (D) mutant ACTB (p.S348L). Yellow dashed line indicates the leading edge of migrating MDCK cells. Scale bar, 50 μm. (E and G) MDCK cells overexpressing either GFP labeled wild-type ACTB or (F and H) mutant ACTB (p.S348L). White arrowheads indicate cell-cell junctions. Scale bar, 10 μm. (I and K) Images of *Xenopus* animal cap cells injected with either wild-type ACTB or (J and L) mutant ACTB mRNA with RFP-CAAX mRNA to label cell-cell junctions. Yellow arrowheads indicate the epithelial cell junctions. Scale bar, 50 μm.

### Mutant ACTB (p.S348L) results in failure of localizing at epithelial cell junction

From a detailed observation of MDCK cells that overexpressed ACTB in a previous experiment, we noticed that mutant ACTB (p.S348L) in MDCK cells showed different localization compared with wild-type ACTB. Wild-type ACTB showed a strong signal at the cell junction, which overlapped with phalloidin staining which detects F-actin (Figure 3E and G). In contrast, the mutant ACTB (p.S348L) did not show an intense signal at the cell junction but rather showed uniform expression in the cytoplasm (Figure 3F and H). In both experiments, the luminance value of GFP at the cell junction was significantly lower in cells that overexpressed the mutant ACTB (p.S348L) (Supplemental figure 3B).

To confirm that this phenomenon was also true *in vivo*, we employed animal cap cells of developing *Xenopus leavis* embryos, which were overexpressed with mutant ACTB (p.S348L). Wild-type ACTB showed a strong signal in the epithelial cell junction, similar to that in MDCK cells (Figure 3I and K). Interestingly, the mutant ACTB (p.S348L) in *Xenopus leavis* animal cap cells failed to localize to the cell-cell junction (Figure 3J and L). These results indicate that the variant detected in the present BWCFF case inhibited ACTB from localizing to the epithelial cell-cell junction.

### Effect of mutated ACTB (p.S348L) on actin dynamics

The dynamic behavior of actin molecules, encompassing both the polymerization and depolymerization processes, plays a pivotal role in modulating the homeostatic equilibrium of the actin complex, which is critical for various cellular activities[15]. For example, actin proteins assemble into fibrous actin filaments that are crucial for cell adhesion and migration[5, 6]. Latrunculin A compound inhibits actin polymerization through its binding affinity to the ATP-binding domain of actin proteins[3, 16]. In the present study, we observed cell surface shrinkage and fragmentation of GFP-labeled wild-type ACTB at the edges of MDCK cells after treatment with Latrunculin A (Figure 4A and B). The cells administered mutant ACTB (p.S348L) also showed shrinkage after treatment with Latrunculin A, whereas GFP-labeled ACTB did not show a fragmented appearance (Figure 4D and E). Fixed cells were stained with phalloidin to detect F-actin. Multiple fragmented yellow dots with overlapping phalloidin (red) and GFP-labeled wild-type ACTB (green) were observed (Figure 4C). In contrast, MDCK cells overexpressing the mutant ACTB (p.S348L) did not show fragmented GFP, whereas phalloidin exhibited fragmentation similar to that of cells overexpressing wild-type ACTB (Figure 4F). These results indicate overexpressed mutant ACTB (p.S348L) have not incorporated in endogenous F-actin which is marked by Phalloidin thus did not change the behavior by the treatment of actin polymerization inhibitor Latrinculin.

**Figure 4.**
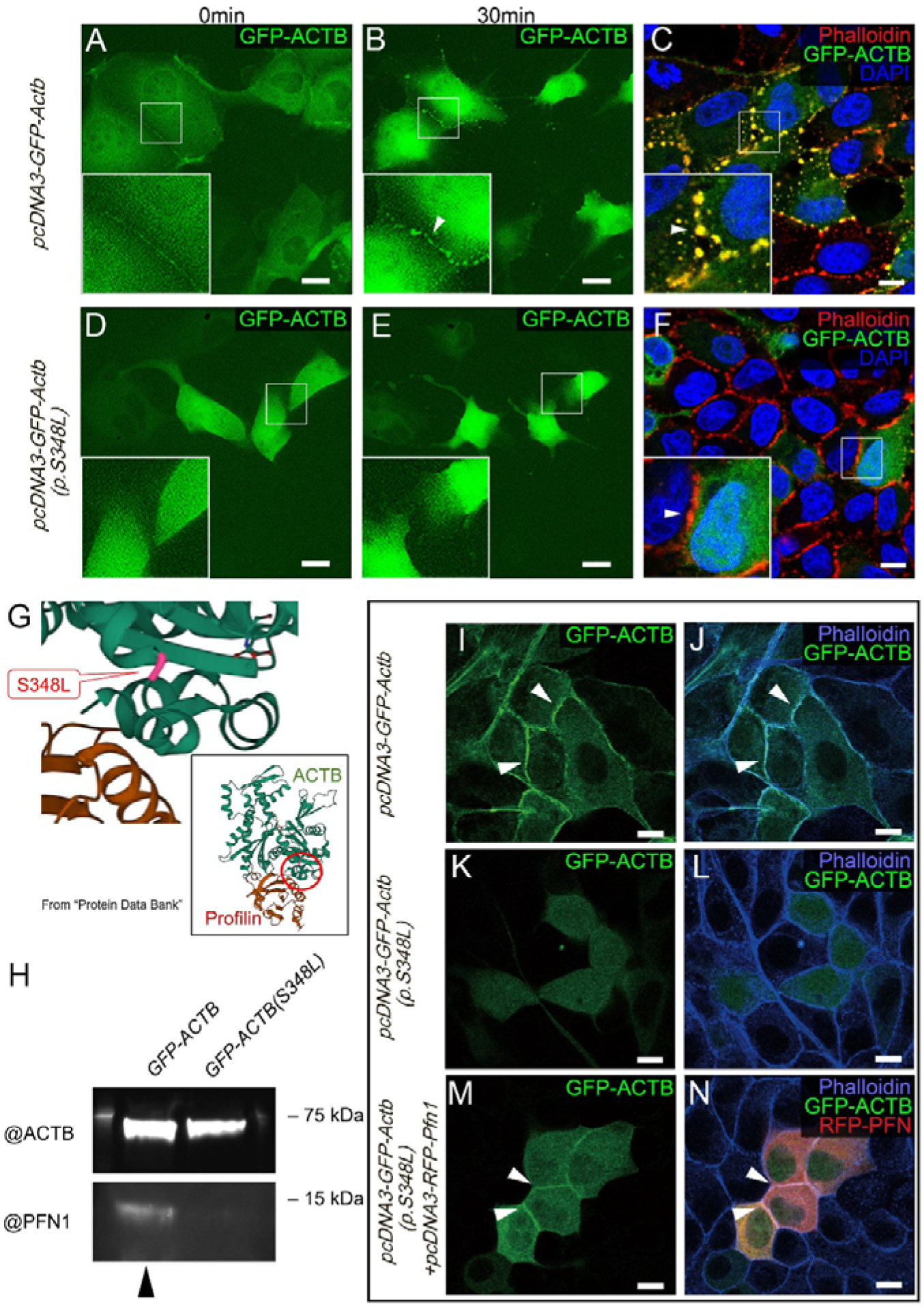
The effect of mutant actin beta (ACTB) (p.S348L) for actin dynamics. (A, B, D, and E) Live imaging of MDCK cells which were transfected with GFP labeled wild type ACTB (A) and mutant ACTB (p.S348L) (D) which was immediately after treated with Latrunclin A. (B and E) Same cells with A and D after 30 min of Latrunculin A treatment. (C) Fixed MDCK cells transfected with wild type ACTB and (D) mutant ACTB (p.S348L) treated with Latrunculin A. White arrowheads indicate cell surface shrinkage as well as fragmentation of GFP-labeled ACTB at the cell-cell junction. Scale bar, 10 μm. (G) Three-dimensional structure of ACTB-PFN1 complex image from the RCSB PDB (https://www.rcsb.org/structure/2btf). The red circle indicates the position of ACTB (p.S348L). (H) Immunoprecipitation of GFP-labeled ACTB from the mixture of exogeneous PFN1 and cellular extract in MDCK cells transfected with wild type and mutant ACTB (p.S348L). MDCK cells transfected with wild type ACTB (I and J) and mutant ACTB (p.S348L) (K and L). Co-transfection of mutant ACTB (p.S348L) and PFN1 recovered the localization of mutant ACTB (p.S348L) into cell-cell junction (M and N). Scale bar, 10 μm.

### Mutant ACTB (p.S348L) reduces its binding affinity with PROFILIN1 (PFN1)

The dynamics of the actin complexes are supported by various actin-associated proteins. From the structure in the protein database (RCSB.org)(https://www.rcsb.org/structure/2btf)(PDB ID:2BTF), the ACTB variant (p.S348L) was located adjacent to the PFN1 binding site[17] (Figure 4G). Therefore, we investigated the role of PFN1, an actin-binding protein that binds to ACTB monomer and facilitates ADP-ACTB nucleotide exchange, leading to ATP-ACTB storage and induction of polymerization into actin filaments. Immunoprecipitation by using the antibody against GFP was performed using the cellular extract of both wild-type and mutant ACTB (p.S348L)-overexpressing MDCK cells. Immunoprecipitated GFP-labeled ACTB was incubated with exogenous PFN1 and subjected to western blot analysis. We detected similar expression levels between mutant (p.S348L) and wild type ACTB, while the signal of PFN1 noticeably reduced in the sample with mutant ACTB (p.S348L) (Figure 4H). These results suggest that the variant compromises the ACTB protein’s affinity for the PFN1 protein which in turn could lead to inhibition of actin polymerization. To test this hypothesis, we overexpressed RFP-labeled PFN1 in conjunction with the mutant ACTB (p.S348L) to evaluate the behavior of mutant ACTB. As previously mentioned, overexpressed wild-type ACTB exists at the cell junction with phalloidin (Figure 4I and J), whereas mutant ACTB (p.S348L) lacks this ability (Figure 4K and L). Interestingly, PFN1 overexpression partially restored the localization of mutated ACTB (p.S348L) to the cell junction (Figure 4M and N). These findings imply that the presence of the mutated ACTB (p.S348L) reduces its binding affinity to PFN1, resulting in a deviant behavior.

### Genome editing of *ACTB* in *Xenopus* embryos resulted in craniofacial phenotype

To further investigate the influence of mutant ACTB (p.S348L) on craniofacial development, genome editing with CRISPR/Cas9 was performed on *X. laevis* embryos. The sequence of the guide RNA to two *ACTB* homologs, *actb.L* and *actb.S* is described in green, which could make the double-strand break (DSB, yellow arrowhead) approximate S348, as indicated in red (Figure 5A). Compared with the linear shape of mouth opening in the uninjected (Figure 5B) and *tyr* F0 mutants (Figure 5C), the experimental *actbL/S* crispants exhibited a substantial increase in embryos manifesting craniofacial deformities, including facial clefts which is shown by small mouth opening and obvious notch at the upper jaw (Figure 5D). Additionally, the intercanthal distance and mouth width were measured in each group (Figure 5E). We detected a significant reduction in both lengths in the *actb.L/S* crispants group, while no significant difference was observed between the uninjected and *tyr.L/S* crispants groups (Figures 5F and G). These results strongly indicated the impact of mutated *ACTB* during embryonic craniofacial development *in vivo*.

**Figure 5.**
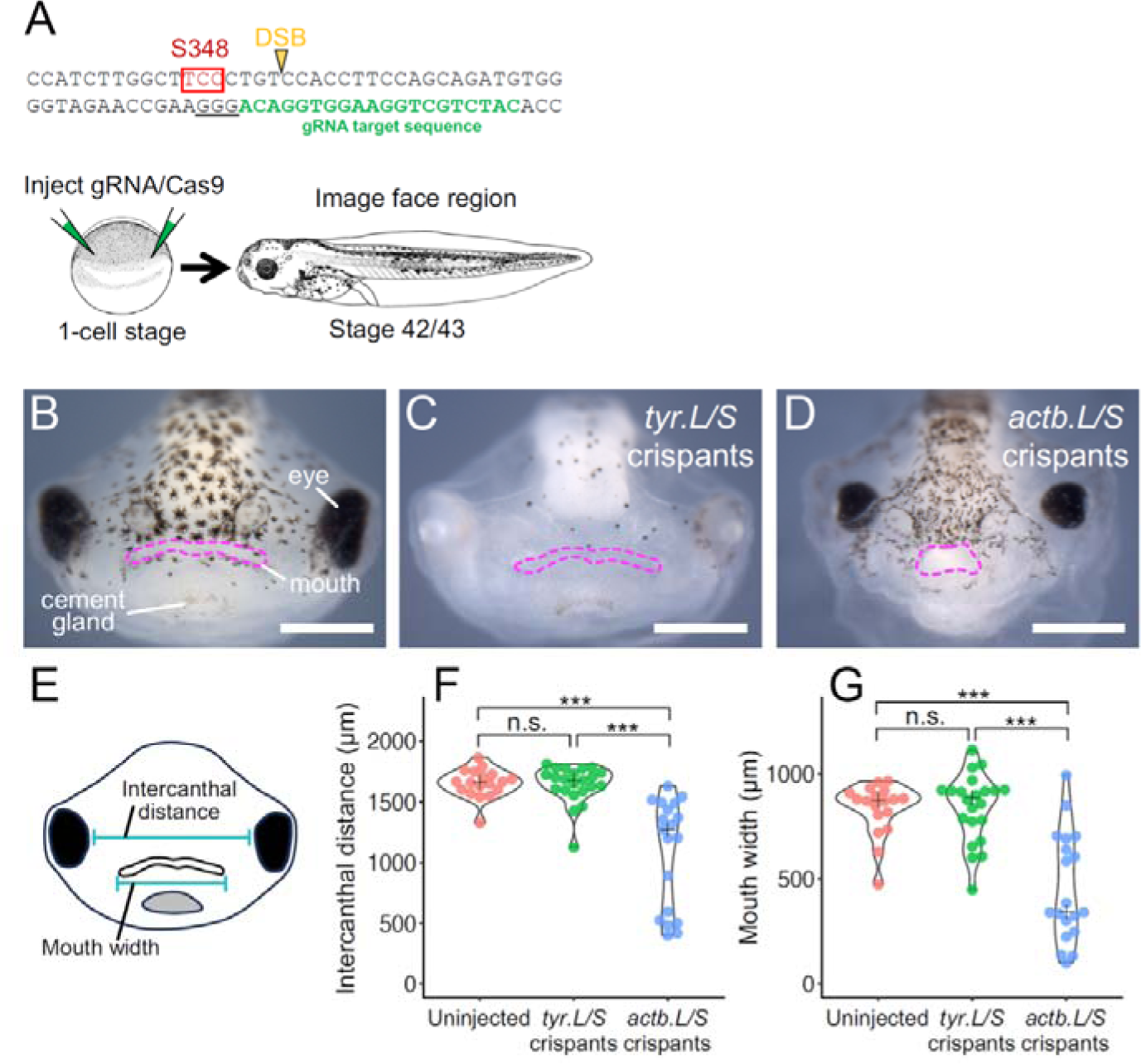
Craniofacial defects induced by disrupting *actb* in *Xenopus*. (A) Schematic drawing of the locus of *actb* S348(red), guide RNA (green) and the predicted double strand break (yellow). The sequences of *actb.L* and *actb.S* in this region are identical. (B) Frontal view of *Xenopus* face of uninjected and (C) *tyr.L/S* crispants and (D) *actb.L/S* crispants. (E) Schematic drawing showing the measurements for intercanthal distance and mouth width. Violin plots of intercanthal distance (F) and mouth width (G) across three groups. An asterisk indicates statistical significance (***P<0.001).

## Discussion

The spectrum of diseases associated with the mutated actin family is defined as actinopathies, which is characterized by a wide range of systemic phenotypes depending on the class of actin that harbors a variant[1]. ACTB and actin gamma are non-muscle actin proteins that are the major components of the cytoskeleton. Thus, their variants have a wide range of effects throughout the body, such as BWCFF [18]. The BWCFF case presented here exhibited *de novo* missense variant in Exon5 of *ACTB* (p.S348L) with multiple craniofacial phenotypes, including cleft lip and palate.

ACTB is a housekeeping protein that is ubiquitously expressed in most cellular entities within an organism. Notably, based on our expression analysis, *ACTB* exhibited widespread distribution, with certain tissues demonstrating heightened expression in the mouse embryo, specifically in the maxillary complex at E10.5, or the epithelial seam during secondary palate fusion at E14.5 (Figure 1). ACTB functions as a major component of stress fibers in cell-cell junctions and the peripheral edges of cells, depending on the cellular subtype and state[19]. These results suggest that different cells require ACTB at varying levels, as certain cells display a greater intensity than others. This divergence in tissue-specific and status-dependent ACTB levels may underpin the mechanism governing the disparate phenotypic degrees observed in BWCFF across distinct tissues.

Based on these results and the significance of epithelial cell behavior in palatal fusion, we further investigated the biological importance of ACTB (p.S348L) in an epithelial cell line (MDCK). MDCK cells are commonly employed to model epithelial cell behavior, including migration during wound healing, and are occasionally used to simulate palatal fusion[20]. Interestingly, the group of cells overexpressing ACTB (p.S348L) showed slower wound closure than cells that overexpressed wild-type ACTB (Figure 3A-D). The epithelial tissue between the fusing facial prominences must be removed via several mechanisms. Among these mechanisms, epithelial cell migration governed by actomyosin contractility is critical for palatal fusion, and its retardation results in facial clefts[9, 12]. Additionally, epithelial cells have been shown to require proper actin polymerization and lamellipodia formation for migration[21]. These results suggest that overexpression of mutant ACTB (p.S348L) slows down epithelial cell migration, which could influence the behavior of embryonic palatal epithelium. Moreover, we observed a diminished signal for the overexpressed mutant ACTB (p.S348L) in contrast to the wild-type ACTB at the intercellular junctions of MDCK cells. Remarkably, this phenomenon was reproduced in animal cap cells of *Xenopus* embryos (Figure 3I-L). Animal cap of Xenopus left *in situ* would eventually develop into ectodermal tissue and thus used for investigating the role of exogeneous proteins in ectodermal cells. Especially, combination of animal cap and MDCK cells was used for modeling embryonic epithelial behaviors and molecular dynamics on cell-cell junction[22]. In these set of experiments, we revealed good evidence of ACTB (p.S348L) lose the ability to localize on the epithelial cell-cell junction both *in vitro* and *vivo*. Actin polymerization is also critical for the proper localization of actin bundles in cells, such as stress fibers and cell-cell junctions[23]. During the growth and fusion of the embryonic facial prominence, surface ectoderm cells must contact the opposing palatal shelf, which requires epithelial adhesion. Notably, variants in the E-cadherin–P120 catenin complex, a vital constituent of adherens junctions that are crucial for epithelial adhesion, have been associated with familial facial clefts in humans[24]. Furthermore, the polymerized actin complex plays a critical role in epithelial adherens junctions by interacting with various classes of cadherins and catenin proteins[25]. The limited capability of ACTB(p.S348L) to localize to cell-cell junctions in epithelial cells observed in this study may reflect a defect in actin polymerization, consequently impairing surface epithelial cell adhesion in the affected individual and contributing to the orofacial clefting phenotype.

To further investigate the mechanism underlying the effect for retarded cell migration and mislocalization, we focused on PFN1, a protein critical for actin polymerization that has been shown to bind to the adjacent area of ACTB (p.S348L)[15, 26]. Immunoprecipitation showed that wild-type ACTB effectively precipitated PFN1, whereas the quantities were noticeably diminished by mutant ACTB (p.S348L) in MDCK cells. Notably, the mutant ACTB (p.S348L) regained its ability to localize at the cell-cell junction when PFN1 was overexpressed simultaneously (Figure 4H-N). These results indicated that the reduced ability of mutant ACTB(p.S348L) to bind PFN1 underlies the mechanism of malfunctioned protein. A missense variant could be associated with protein malfunction due to misfolding, altering the affinity with associating proteins, in this case, possibly PFN1.

Inhibition of actin polymerization using Latrinculin-A in ACTB overexpressed cells also revealed noticeable actin fragmentation with wild-type ACTB but not with mutant ACTB (p.S348L) (Figure 4A-F). These results indicate the effect of p.S348L on the behavior of ACTB. First, there is a possibility that actin polymerization is retarded, as actin polymerization is required for proper F-actin localization, such as in cell-cell junctions. Second, it is also possible to think that the p.S348L variant makes F-actin resistant to disassembly. From the set of results, mutant ACTB (p.S348L) shows a reduced ability to bind PFN1, supporting the likelihood of the first theory. However, further study is essential to prove the effect of p.S348L on actin polymerization.

We conducted a functional assay employing CRISPR/Cas9 to perturb the *Actb* gene, with the guide RNA strategically inducing a double-stranded break around serine (S) at the 348th position of *actb* in developing *Xenopus* embryos, resulting in the manifestation of craniofacial defects (Figure 5). This outcome strongly implies that ACTB (p.S348L) exerted a considerable influence on the etiology of craniofacial defects in the present case of BWCFF.

Previous studies have investigated the role of ACTB (p.S348L)[13, 14]. Several common human phenotypes encompassing craniofacial anomalies have been observed in these reports. However, other studies did not explicitly describe the presence of any form of orofacial cleft in these patients. There are several possible explanations for this phenotypic variation within the same variant in patients with *BWCFF*. First, subtle manifestations of orofacial cleft may have been disregarded. In particular, microcleft or submucous cleft is sometimes challenging to detect in newborns, and some patients remain undiagnosed. Second, the prevalence of orofacial cleft could be influenced by genetic background; the East Asian population exhibits a higher incidence of orofacial cleft than other populations, which influences the phenotypic variation among these BWCFF patients[1, 3, 27]. There is speculation regarding East Asian specificity in exhibiting certain genetic factors as modifiers that increase the incidence of orofacial clefts, but this requires further confirmation. Investigating the relationship between these race-specific genetic influences and ACTB in future studies would be intriguing.

Functional assays involving different variants in *ACTB* and employing various models have been proposed, with certain variants demonstrating a gain-of-function in specific cell types, including lymphoblasts and yeast cells[3, 13, 28]. To the best of our knowledge, this study is the first to assess the biological implications of *ACTB* variants associated with BWCFF in epithelial tissue and orofacial cleft development. Additionally, there have been proposals positing a defect in cranial neural crest cell development due to an *ACTB* variant based on both the craniofacial phenotype of BWCFF and animal studies[1, 29]. *Actb* null embryos exhibit elevated cell death in pre-migratory neural crest cells which could be one of the underlying mechanism for broad craniofacial phenotype[29]. For these reasons, it is also reasonable to surmise that ACTB (p.S348L) profoundly influences cranial neural crest cell development, synergistically leading to a myriad of craniofacial defects.

## Materials and Methods

### Molecular analysis

In collaboration with the Initiative on Rare and Undiagnosed Diseases, we conducted a trio exome analysis of a family presenting with a patient exhibiting systemic phenotypes, including bilateral cleft lip and palate. Informed consent was obtained from the patients and their parents in accordance with the institutional review board. Whole exome sequencing was performed on genomic DNA extracted from the peripheral lymphocytes of the patients and their families using the SureSelect Human All Exon Kit V6 (Agilent Technologies, Santa Clara, CA, USA) with sequencing on the NovaSeq 6000 platform (Illumina, San Diego, CA, USA). We checked the quality of the FASTQ files using FASTQC (https://www.bioinformatics.babraham.ac.uk/projects/fastqc/) and removed low-quality reads using trimmomatic-0.36 (http://www.usadellab.org/cms/?page=trimmomatic). Quality-checked reads were aligned to GRCh37 using the Burrows-Wheeler Aligner (http://bio-bwa.sourceforge.net/) and variants were called using the GATK HaplotypeCaller. These genes were annotated using ANNOVAR (http://annovar.openbioinformatics.org/en/latest/). In silico analyses of the variants were performed using CADD (http://cadd.gs.washington.edu/) and PROVEAN (http://provean.jcvi.org/seq_submit.php). High-frequency (minor allele frequency > 5% in the Japanese population), synonymous, and intergenic variants were manually filtered out.

### *In situ* hybridization, immunohistochemistry and immunocytochemistry

Wild-type pregnant ICR mice were purchased from CLEA Japan, Inc. Dissected embryos were fixed in 4% Paraformaldehyde for overnight at 4lJ. Immunohistochemistry was performed using the M.O.M. Immunodetection Kit (VECTOR), in accordance with the manufacturer’s protocol. Briefly, 12-μm thick frozen sections were incubated with selected primary antibodies that recognize ACTB (Proteintech) and E-cadherin (Cell Signaling Technology) after heat-induced antigen retrieval. The samples were subsequently incubated with appropriate secondary antibodies for 1 h at 37lJ and counterstained with DAPI (; Sigma-Aldrich). Cultured cells were fixed with 4% Paraformaldehyde for 15 min, followed by permeabilization with 0.1% Triton X-100 for 5 min. The samples were counterstained with Alexa Fluor 594 or 350 phalloidin (Invitrogen; 1:250) for 1 h and DAPI (Sigma-Aldrich) (1:1,000) for 20 min to visualize F-actin and nuclei. The samples were mounted with a fluorescent mounting medium (DakoCytomation) and visualized using either a Leica TCS SP8 (Leica) or BZ-X710 All-in-one Fluorescence Microscope (Keyence). *In situ* hybridization was performed as previously described[30]. The primer sets used to produce the antisense oligo probe of mouse *Actb* were as follows: sense 5-CACACCTTCTACAATGAGCTGC −3 and antisense 5-GGCATAGAGGTCTTTACGGATG −3 according to the Allen Brain Atlas (https://atlas.brain-map.org/). Following *in situ* hybridization, heads were embedded in Tissue-Tek (OCT compound, Sakura), and cut into 20-μm-thick frozen sections for histological observation.

### Plasmid construction

The nucleotide sequences encoding *ACTB* and its mutated form (MANE Select *ACTB*(NM_001101.5):c.1043C>T (p.Ser348Leu) GRCh37 Chr7:5567464) and *PFN1* were amplified and cloned into the pcDNA3-GFP vector, resulting in the production of a fusion protein of ACTB and PFN1 with EGFP and RFP, respectively, under the control of the CMV promoter. This was performed using the In-Fusion HD Cloning Plus Kit (Takara Bio, Shiga, Japan), following the manufacturer’s instructions. For *Xenopus*, GFP-fused Actb WT and S348L in pcDNA3-GFP were amplified using KOD One (TOYOBO, Osaka, Japan) and cloned into pCSf107mT[31] using the In-Fusion HD Cloning Kit (Takara, Shiga, Japan) resulting in pCSf107mT-GFP-Actb WT and S348L. mCherry was amplified and cloned into pCS2p to generate pCS2p-mCherry. The sequences of these plasmids were confirmed by Sanger Sequencing. pCS2-mRFP-CAXX (membrane-targeted mRFP) has been described previously[32].

### mRNA preparation

Capped mRNAs for microinjection into *Xenopus* embryos were synthesized using an mMessage mMachine SP6 Transcription Kit (AM1340; Thermo Fisher Scientific, MA, USA) and purified by phenol/chloroform extraction using an RNeasy Mini Kit (QIAGEN, Venlo, Netherlands).

### Guide RNA and Cas9 protein preparation

A crRNA (5’-CATCTGCTGGAAGGTGGACA-3’) was designed to target the exon 6 of *X. laevis actb.L* and *actb.S* genes with *Xenopus* genome database (http://viewer.shigen.info/xenopus/) and CHOPCHOP[33]. crRNA, tracrRNA (#1072533), and Cas9 protein (#1081059) were purchased from IDT (Coralville, MD, USA). The crRNA-tracrRNA duplex was formed as previously described[34]. The background effect of genome editing was assessed by designing guide RNA for two tyrosinase homologs, *tyr.L* and *tyr.S* as previously described[35, 36]. The Cas9 protein and gRNAs were incubated at 37lJ for 5 min in a buffer containing 150 mM KCl and 20 mM HEPES (pH 7.5) to form Cas9-gRNA RNP complexes, then kept at room temperature until microinjection.

### Embryo manipulation and microinjection of *Xenopus*

Wild-type *X. laevis* adults were purchased from Watanabe Zoushoku (Hyogo, Japan) and kept in a recirculating aquarium system with a water temperature of 20lJ. *In vitro* fertilization, dejellying, and embryo staging were performed as previously described[37, 38]. For mRNA overexpression, 200 pg GFP-Actb WT or S348L mRNA and 250 pg RFP-CAAX mRNA were injected into the animal pole of two-cell stage embryos in 3% Ficoll/0.3 × Marc’s Modified Ringer’s medium (MMR). The injected embryos were cultured in the same medium until they reached stage 9.

For the Cas9-RNP injection, 20 nL of the injection mixture containing 300 pg gRNA, 4 ng Cas9 protein, and 100 pg mCherry mRNA was injected radially (10 nL × 2) into the marginal zone of one-cell-stage embryos in 3% Ficoll/0.3 × MMR within 40 min after fertilization. The injected embryos were kept in the same medium at 22lJ until the following day, then cultured in 0.3 × MMR until appropriate stages at 22lJ.

### Genotyping

Tailbud stage embryos with uniform mCherry fluorescence were treated at 95lJ for 5 min in the lysis buffer (10 mM Tris-HCl pH 8, 1 mM EDTA, 0.1% Nonidet P-40), then digested with Proteinase K (Takara) at 55lJ overnight. Heat-inactivated supernatants were collected after centrifugation of crude genomic DNA samples. The 324-bp *actb.L* target region was amplified from crude DNA samples using KOD FX Neo (TOYOBO). The amplified PCR products were cleaned using the EnzSAP PCR Clean-Up Reagent Kit (Edge BioSystems, CA, USA) and analyzed by direct sequencing. The sequences of PCR primers used are as follows: forward primer, 5’-GCACCATGAAAATCAAGGTATG-3’; reverse primer, 5’-AAGAGTAAAGCCATGCCAATGT-3.’ The efficiency of indel formation was analyzed using the Inference of CRISPR Edits tool[39].

### Imaging of *Xenopus* embryos

Stage 9–10 embryos were mounted in 1% low-melting agarose/0.3 × MMR on a glass-based dish (3910-035; AGC, Shizuoka, Japan). Confocal fluorescence images were acquired using FV1000-D (Olympus, Tokyo, Japan) with a 40× (UPLSAPO 40x NA0.90; Olympus) objective lens at a room temperature adjusted to 18–20lJ. Bright-field images were acquired using an Axio Zoom.V16 zoom microscope (Zeiss, Oberkochen, Germany), and analyzed using ZEN (Zeiss) and Fiji software. The violin plot was created using the ggplot package in RStudio based on the measurements of intercanthal distance and mouth width as it was described [40].

### Cell lines and cell culture

MDCK cells (JCRB cell bank: IFO50071) were cultured in _α_MEM with nucleosides (Gibco) and 10% FBS (Gibco) under standard conditions. For electroporation, MDCK cells were transfected with overexpression vectors containing either wild-type or mutant (c.1043C > T) ACTB in Opti-MEM (Gibco) using a NEPA21 electroporator (Nepagene). Five square pulses of 50-ms duration with 50-ms intervals at 20 V were applied.

For migration assays, MDCK cells transfected with either construct seeded onto fibronectin-coated 35 mm glass bottom dishes (Matsunami Glass Ind., Ltd) in a two-well silicone insert with a defined cell-free gap (Ibidi). After the cells reached confluence at 24 h, the culture insert was removed, and the area of cell movement was quantified 10 h after removal of the silicone insert.

MDCK cells transfected with either construct were treated with Latrunculin A (AdipoGen) at a concentration of 100uM. Live imaging was performed using a BZ-X710 All-in-one Fluorescence Microscope (Keyence, Osaka, Japan).

### Immunoprecipitation and Western Blot

Sample protein was extract from MDCK cells which was transfected with GFP-tagged wild-type ACTB and mutant ACTB (p.S348L) which was mixed with the recombinant PFN1 to let them bind to extracted protein with concentration of 2 µM. The GFP-Trap agarose kit (Chromotek) was used for immunoprecipitation according to the manufacturer’s protocol. The pulled down proteins were separated by Sodium dodecyl sulfate Poly-Acrylamide Gel Electrophoresis (SDS-PAGE) and transferred to polyvinylidene difluoride membranes (Bio-Rad Laboratories). Membranes were incubated with anti-ACTB (Proteintech) and anti-PFN1 (Santa Cruz Biotechnology) antibodies. The bound antibodies were detected using horseradish peroxidase-conjugated antibodies (Cell Signaling Technology) and an ECL detection kit (Bio-Rad Laboratories) according to the manufacturer’s protocol.

### Statistics

Mann-Whitney test with Bonferroni adjustment was used in Figure 5. Two-tailed Student’s *t*-tests were performed on the data presented in Supplemental figure 3. P was set at p < 0.05 in all experiments.

## Data availability

The data that support the findings of this study are not publicly available, but are available from the corresponding author upon reasonable request.

## Ethics declaration

Written informed consent was obtained from the patient for the publication of their photo. The study was reviewed and approved by the Committee on the Ethics of Animal Experiments of Osaka University Graduate School of Dentistry and the Ethics Committee of Osaka University.

## Acknowledgement

This study was supported in part by grants from the Initiative on Rare and Undiagnosed Diseases project of the Japanese Agency for Medical and Development (JP23bk0304002), Ministry of Education, Science, Sports, and Culture of Japan (no. 19H03858 to HK), the NOVARTIS Foundation (Japan) for the Promotion of Science and Takeda Science Foundation. We would like to thank the National BioResource Project (Clawed Frogs/Newts) of MEXT at the Hiroshima University Amphibian Research Center (RRID: SCR_019015) for their technical assistance. We thank Dr. Shoichiro Ono for valuable insights into conducting this study. We also wish to express our sincere appreciation to the patients and their families for their consent to participate in this study.

---

**Supplemental figure 1.**
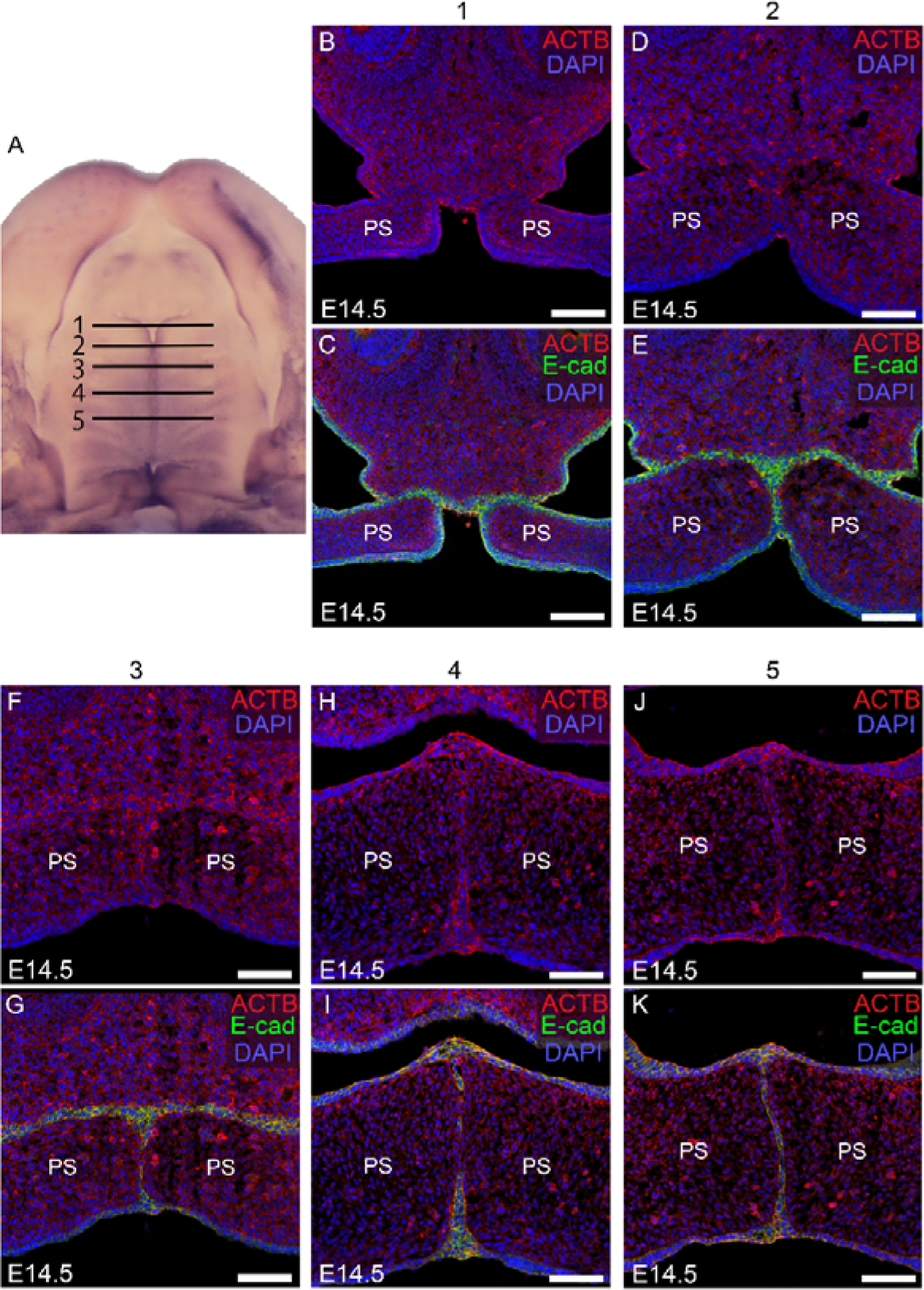
Expression of ACTB during palatogenesis. (A) Ventral view of dissected maxilla at E14.5. Frontal sections of E14.5 maxilla at the locations of black lines shown in B-K. (B-K) Immunohistochemistry of ACTB (red) and E-cadherin (green) of frontal section of embryonic head of E14.5. PS, palatal shelf. Scale bar, 50 μm.

**Supplemental figure 2.**
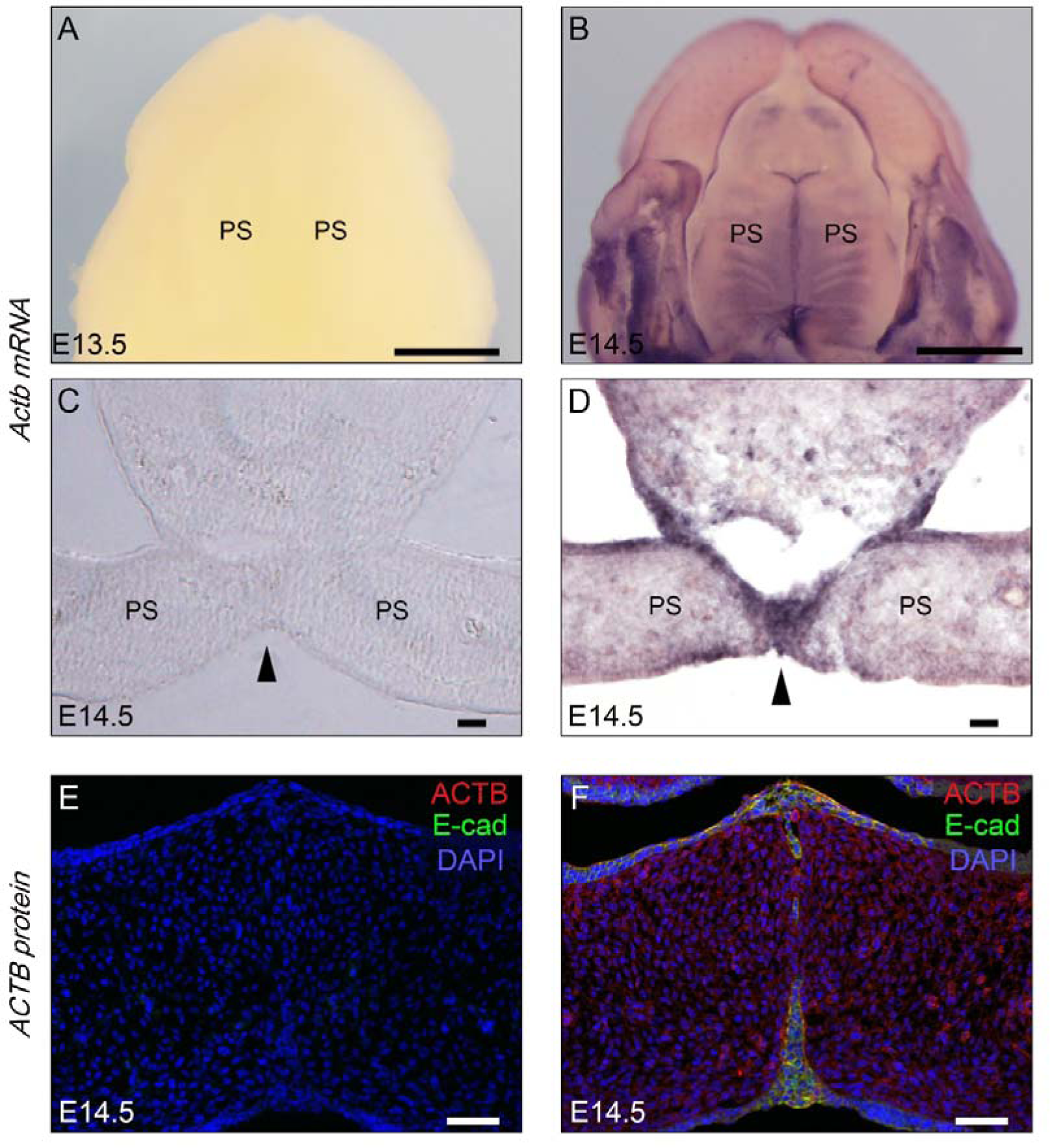
Negative controls of *In situ*hybridization and immunohistochemistry. (A, C and E) negative controls of each B, D and F, which corresponds to C, D, and F in Fig. 2. PS, palatal shelf. Scale bar, 1 mm (A–D) and 50 μm (E, F).

**Supplemental figure 3.**
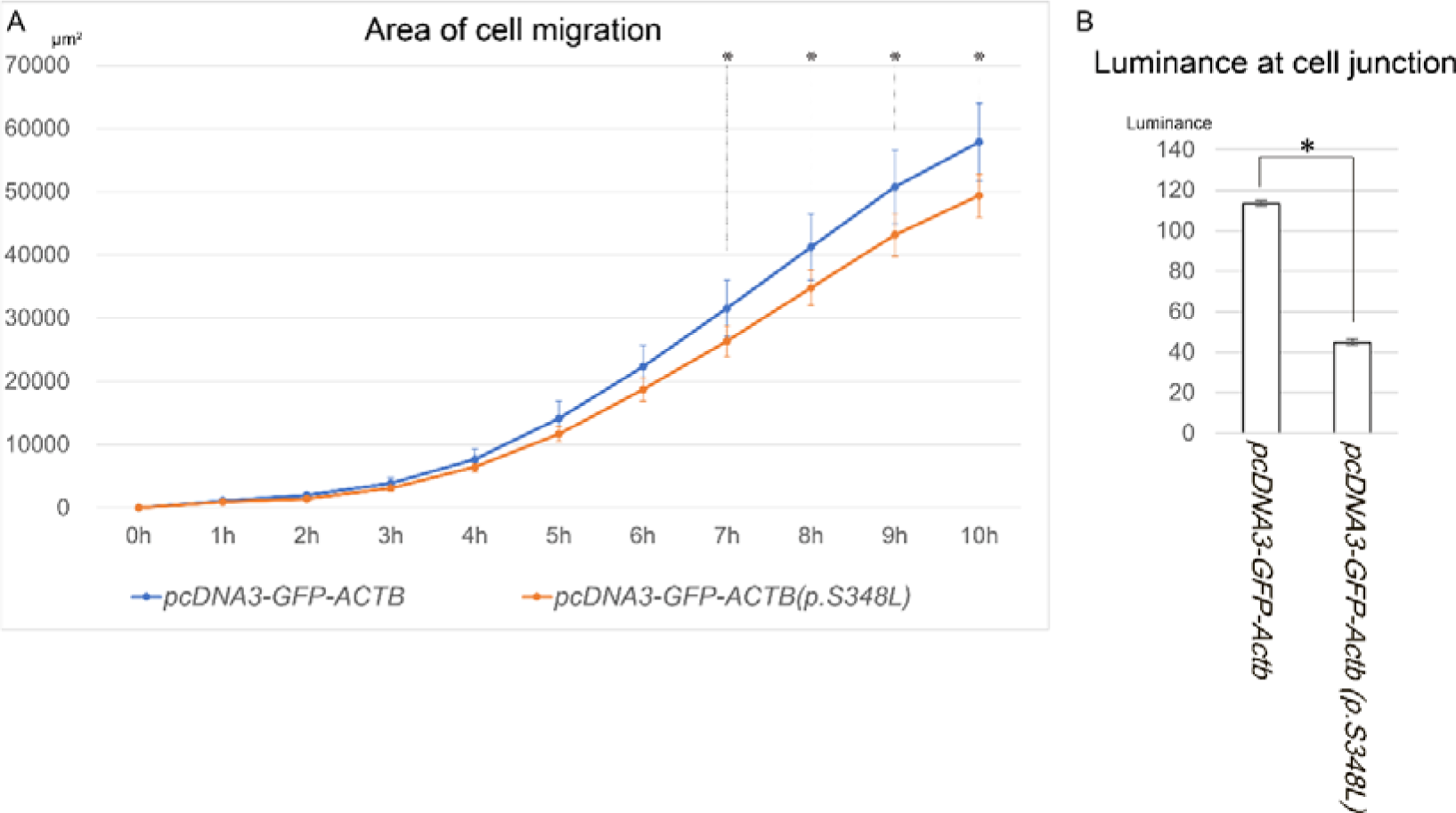
Area of cell migration and luminance of GFP in cell junction. (A) The cell migration area depicted in Figure 3A-D, measured over time through three experimental repetitions. The blue and orange lines represent the populations of wild-type ACTB and mutant ACTB (p.S348L) overexpressing cells, respectively. *p<0.05. (B) The luminance values of GFP at cell-cell junctions for both wild-type ACTB and mutant ACTB (p.S348L), presented in Figure 3E-H. *p<0.05.

## REFERENCES

1. Verloes A, Di Donato N, Masliah-Planchon J, Jongmans M, Abdul-Raman OA, Albrecht B, et al. Baraitser-Winter cerebrofrontofacial syndrome: delineation of the spectrum in 42 cases. Eur J Hum Genet. 2015;23(3):292–301. Epub 2014/07/24. doi: 10.1038/ejhg.2014.95. PubMed PMID: 25052316; PubMed Central PMCID: PMCPMC4326722.

2. Baraitser M, Winter RM. Iris coloboma, ptosis, hypertelorism, and mental retardation: a new syndrome. J Med Genet. 1988;25(1):41–3. doi: 10.1136/jmg.25.1.41. PubMed PMID: 3351890; PubMed Central PMCID: PMCPMC1015421.

3. Rivière JB, van Bon BW, Hoischen A, Kholmanskikh SS, O’Roak BJ, Gilissen C, et al. De novo mutations in the actin genes ACTB and ACTG1 cause Baraitser-Winter syndrome. Nat Genet. 2012;44(4):440–4, S1-2. Epub 20120226. doi: 10.1038/ng.1091. PubMed PMID: 22366783; PubMed Central PMCID: PMCPMC3677859.

4. Dominguez R, Holmes KC. Actin structure and function. Annu Rev Biophys. 2011;40:169-86. doi: 10.1146/annurev-biophys-042910-155359. PubMed PMID: 21314430; PubMed Central PMCID: PMCPMC3130349.

5. Braga V. Spatial integration of E-cadherin adhesion, signalling and the epithelial cytoskeleton. Curr Opin Cell Biol. 2016;42:138–45. Epub 20160806. doi: 10.1016/j.ceb.2016.07.006. PubMed PMID: 27504601.

6. Vaezi A, Bauer C, Vasioukhin V, Fuchs E. Actin cable dynamics and Rho/Rock orchestrate a polarized cytoskeletal architecture in the early steps of assembling a stratified epithelium. Dev Cell. 2002;3(3):367–81. PubMed PMID: 12361600.

7. Trepat X, Chen Z, Jacobson K. Cell migration. Compr Physiol. 2012;2(4):2369-92. doi: 10.1002/cphy.c110012. PubMed PMID: 23720251; PubMed Central PMCID: PMCPMC4457291.

8. Bush JO, Jiang R. Palatogenesis: morphogenetic and molecular mechanisms of secondary palate development. Development. 2012;139(2):231–43. doi: 10.1242/dev.067082. PubMed PMID: 22186724; PubMed Central PMCID: PMCPMC3243091.

9. Aoyama G, Kurosaka H, Oka A, Nakatsugawa K, Yamamoto S, Sarper SE, et al. Observation of Dynamic Cellular Migration of the Medial Edge Epithelium of the Palatal Shelf. Front Physiol. 2019;10:698. Epub 20190606. doi: 10.3389/fphys.2019.00698. PubMed PMID: 31244674; PubMed Central PMCID: PMCPMC6562562.

10. Cuervo R, Valencia C, Chandraratna RA, Covarrubias L. Programmed cell death is required for palate shelf fusion and is regulated by retinoic acid. Dev Biol. 2002;245(1):145–56. doi: 10.1006/dbio.2002.0620. PubMed PMID: 11969262.

11. Shuler CF, Halpern DE, Guo Y, Sank AC. Medial edge epithelium fate traced by cell lineage analysis during epithelial-mesenchymal transformation in vivo. Dev Biol. 1992;154(2):318–30. doi: 10.1016/0012-1606(92)90071-n. PubMed PMID: 1385235.

12. Kim S, Lewis AE, Singh V, Ma X, Adelstein R, Bush JO. Convergence and extrusion are required for normal fusion of the mammalian secondary palate. PLoS Biol. 2015;13(4):e1002122. Epub 20150407. doi: 10.1371/journal.pbio.1002122. PubMed PMID: 25848986; PubMed Central PMCID: PMCPMC4388528.

13. Sibbin K, Yap P, Nyaga D, Heller R, Evans S, Strachan K, et al. A de novo ACTB gene pathogenic variant in identical twins with phenotypic variation for hydrops and jejunal atresia. Am J Med Genet A. 2022;188(4):1299–306. Epub 20211231. doi: 10.1002/ajmg.a.62631. PubMed PMID: 34970864; PubMed Central PMCID: PMCPMC9302691.

14. Fakhro KA, Robay A, Rodrigues-Flores JL, Mezey JG, Al-Shakaki AA, Chidiac O, et al. Point of Care Exome Sequencing Reveals Allelic and Phenotypic Heterogeneity Underlying Mendelian disease in Qatar. Hum Mol Genet. 2019;28(23):3970–81. doi: 10.1093/hmg/ddz134. PubMed PMID: 31625567.

15. Ono S. The role of cyclase-associated protein in regulating actin filament dynamics - more than a monomer-sequestration factor. J Cell Sci. 2013;126(Pt 15):3249–58. doi: 10.1242/jcs.128231. PubMed PMID: 23908377; PubMed Central PMCID: PMCPMC3730240.

16. Morton WM, Ayscough KR, McLaughlin PJ. Latrunculin alters the actin-monomer subunit interface to prevent polymerization. Nat Cell Biol. 2000;2(6):376–8. doi: 10.1038/35014075. PubMed PMID: 10854330.

17. Schutt CE, Myslik JC, Rozycki MD, Goonesekere NC, Lindberg U. The structure of crystalline profilin-beta-actin. Nature. 1993;365(6449):810-6. doi: 10.1038/365810a0. PubMed PMID: 8413665.

18. Greve JN, Schwäbe FV, Pokrant T, Faix J, Di Donato N, Taft MH, et al. Frameshift mutation S368fs in the gene encoding cytoskeletal β-actin leads to ACTB-associated syndromic thrombocytopenia by impairing actin dynamics. Eur J Cell Biol. 2022;101(2):151216. Epub 20220315. doi: 10.1016/j.ejcb.2022.151216. PubMed PMID: 35313204.

19. Dugina V, Zwaenepoel I, Gabbiani G, Clément S, Chaponnier C. Beta and gamma-cytoplasmic actins display distinct distribution and functional diversity. J Cell Sci. 2009;122(Pt 16):2980–8. Epub 20090728. doi: 10.1242/jcs.041970. PubMed PMID: 19638415.

20. Yu W, Zhang Y, Ruest LB, Svoboda KK. Analysis of Snail1 function and regulation by Twist1 in palatal fusion. Front Physiol. 2013;4:12. Epub 20130219. doi: 10.3389/fphys.2013.00012. PubMed PMID: 23424071; PubMed Central PMCID: PMCPMC3575576.

21. Ponti A, Machacek M, Gupton SL, Waterman-Storer CM, Danuser G. Two distinct actin networks drive the protrusion of migrating cells. Science. 2004;305(5691):1782-6. doi: 10.1126/science.1100533. PubMed PMID: 15375270.

22. Haigo SL, Hildebrand JD, Harland RM, Wallingford JB. Shroom induces apical constriction and is required for hingepoint formation during neural tube closure. Curr Biol. 2003;13(24):2125–37. doi: 10.1016/j.cub.2003.11.054. PubMed PMID: 14680628.

23. Vasioukhin V, Bauer C, Yin M, Fuchs E. Directed actin polymerization is the driving force for epithelial cell-cell adhesion. Cell. 2000;100(2):209–19. doi: 10.1016/s0092-8674(00)81559-7. PubMed PMID: 10660044.

24. Cox LL, Cox TC, Moreno Uribe LM, Zhu Y, Richter CT, Nidey N, et al. Mutations in the Epithelial Cadherin-p120-Catenin Complex Cause Mendelian Non-Syndromic Cleft Lip with or without Cleft Palate. Am J Hum Genet. 2018;102(6):1143–57. Epub 20180524. doi: 10.1016/j.ajhg.2018.04.009. PubMed PMID: 29805042; PubMed Central PMCID: PMCPMC5992119.

25. Tsukita S, Nagafuchi A, Yonemura S. Molecular linkage between cadherins and actin filaments in cell-cell adherens junctions. Curr Opin Cell Biol. 1992;4(5):834–9. doi: 10.1016/0955-0674(92)90108-o. PubMed PMID: 1419062.

26. Korenbaum E, Nordberg P, Björkegren-Sjögren C, Schutt CE, Lindberg U, Karlsson R. The role of profilin in actin polymerization and nucleotide exchange. Biochemistry. 1998;37(26):9274–83. doi: 10.1021/bi9803675. PubMed PMID: 9649308.

27. Beaty TH, Marazita ML, Leslie EJ. Genetic factors influencing risk to orofacial clefts: today’s challenges and tomorrow’s opportunities. F1000Res. 2016;5:2800. Epub 20161130. doi: 10.12688/f1000research.9503.1. PubMed PMID: 27990279; PubMed Central PMCID: PMCPMC5133690.

28. Procaccio V, Salazar G, Ono S, Styers ML, Gearing M, Davila A, et al. A mutation of beta -actin that alters depolymerization dynamics is associated with autosomal dominant developmental malformations, deafness, and dystonia. Am J Hum Genet. 2006;78(6):947–60. Epub 20060421. doi: 10.1086/504271. PubMed PMID: 16685646; PubMed Central PMCID: PMCPMC1474101.

29. Tondeleir D, Noelanders R, Bakkali K, Ampe C. Beta-actin is required for proper mouse neural crest ontogeny. PLoS One. 2014;9(1):e85608. Epub 20140107. doi: 10.1371/journal.pone.0085608. PubMed PMID: 24409333; PubMed Central PMCID: PMCPMC3883714.

30. Kurosaka H, Iulianella A, Williams T, Trainor PA. Disrupting hedgehog and WNT signaling interactions promotes cleft lip pathogenesis. J Clin Invest. 2014;124(4):1660–71. Epub 20140303. doi: 10.1172/JCI72688. PubMed PMID: 24590292; PubMed Central PMCID: PMCPMC3973078.

31. Mii Y, Taira M. Secreted Frizzled-related proteins enhance the diffusion of Wnt ligands and expand their signalling range. Development. 2009;136(24):4083–8. Epub 20091111. doi: 10.1242/dev.032524. PubMed PMID: 19906850.

32. Suzuki M, Sato M, Koyama H, Hara Y, Hayashi K, Yasue N, et al. Distinct intracellular Ca2+ dynamics regulate apical constriction and differentially contribute to neural tube closure. Development. 2017;144(7):1307–16. Epub 20170220. doi: 10.1242/dev.141952. PubMed PMID: 28219946.

33. Labun K, Montague TG, Krause M, Torres Cleuren YN, Tjeldnes H, Valen E. CHOPCHOP v3: expanding the CRISPR web toolbox beyond genome editing. Nucleic Acids Res. 2019;47(W1):W171–W4. doi: 10.1093/nar/gkz365. PubMed PMID: 31106371; PubMed Central PMCID: PMCPMC6602426.

34. Hoshijima K, Jurynec MJ, Klatt Shaw D, Jacobi AM, Behlke MA, Grunwald DJ. Highly Efficient CRISPR-Cas9-Based Methods for Generating Deletion Mutations and F0 Embryos that Lack Gene Function in Zebrafish. Dev Cell. 2019;51(5):645–57.e4. Epub 20191107. doi: 10.1016/j.devcel.2019.10.004. PubMed PMID: 31708433; PubMed Central PMCID: PMCPMC6891219.

35. Tanouchi M, Igawa T, Suzuki N, Suzuki M, Hossain N, Ochi H, et al. Optimization of CRISPR/Cas9-mediated gene disruption in Xenopus laevis using a phenotypic image analysis technique. Dev Growth Differ. 2022;64(4):219–25. Epub 20220412. doi: 10.1111/dgd.12778. PubMed PMID: 35338712.

36. Wang F, Shi Z, Cui Y, Guo X, Shi YB, Chen Y. Targeted gene disruption in Xenopus laevis using CRISPR/Cas9. Cell Biosci. 2015;5:15. Epub 20150414. doi: 10.1186/s13578-015-0006-1. PubMed PMID: 25897376; PubMed Central PMCID: PMCPMC4403895.

37. Nieuwkoop PDF, J. Normal Table of Xenopus laevis (Daudin). Amsterdam: North-Holland Publ. Co.; 1967 June 9.

38. Sive HL, Grainger, R. M. & Harland, R. M. Early Development of Xenopus Laevis: A Laboratory Manual. Cold Spring Harbor, New York: Cold Spring Harbor Laboratory Press; 2000.

39. Conant D, Hsiau T, Rossi N, Oki J, Maures T, Waite K, et al. Inference of CRISPR Edits from Sanger Trace Data. CRISPR J. 2022;5(1):123–30. Epub 20220202. doi: 10.1089/crispr.2021.0113. PubMed PMID: 35119294.

40. Wyatt BH, Raymond TO, Lansdon LA, Darbro BW, Murray JC, Manak JR, et al. Using an aquatic model, Xenopus laevis, to uncover the role of chromodomain 1 in craniofacial disorders. Genesis. 2021;59(1-2):e23394. Epub 2020/09/13. doi: 10.1002/dvg.23394. PubMed PMID: 32918369; PubMed Central PMCID: PMCPMC10701884.

